# Identification of Six novel missense single nucleotide polymorphisms in the MAOA gene predisposing to aggressive behavior. Bioinformatics study

**DOI:** 10.1101/2019.12.18.880963

**Authors:** Abdelrahman H. Abdelmoenim, Mujahed I. Mustafa, Naseem S. Murshed, Nosiba S. Omer, Alaa I. Mohammed, Rania A. Abdulmajed, Enas dk. Dawoud, Abdelrafie M. Makhawi

## Abstract

**Background:** An astonishing observation is that aggressive behavior is actually a highly heritable. Recent experimental work and behavior research has linked individual variation in a functional polymorphism of the monoamine oxidase-A gene (MAOA) to the occurrence of anger-driven aggression. Aggressive antisocial and violent behavior has become a regularly debated topic in the scientific community; the impending question is what is the source of aggressive behavior, is it genetic or environmental or is it just an individual choice. This study aims to analyses the SNPs found in MAOA gene and it is possible association to aggressive behavior.

**Method:** Various bioinformatics software (SIFT, PolyPhen-2, PROVEAN, SNAP22, SNP&GO and PMut)is used to analyses the SNPs within the MAOA gene to study the structural and functional implication on the associated protein, which is further clarified using chimera software. Then gene-gene interaction is studied with geneMANIA software. Furthermore, conservation and annotation studies were done through the ConSurf server and Variant Effect Predictor (VEP) respectively.

**Result:** Six missense SNPs were found to affect the structural and functional prospect of MAOA protein.

**Conclusion:** Genetic mutation within MAOA is likely to be associated with aggressive behavior; this will enrich future management and screening possibilities for this behavior.

## Introduction

“Aggression is defined as harmful social interaction with the intention of inflicting damage or other unpleasantness upon another individual. It can happen with or without trigger(1). However, aggressive behavior is considered as an adaptation mechanism not only in humans but in other species as well. Furthermore, sometimes the aggressive response can be an indication of psychopathology especially when it exceeds the normal range (2).

Studies had shown that, Aggressive and antisocial behavior is highly heritable (3).When studying the heritability of aggression and antisocial behaviors the MAOA gene is usually the gene of interest.The effect of MAOA in modulating human behavior has been widely studied. Moreover, associations have been described between several psychiatric illnesses and altered MAO-A function. Even more, antisocial personality disorder (ASPD) and borderline personality disorder (BPD) have been the main focus of many studies regarding the MAO A gene in non-clinical samples (4).

a study done between 2000 and 2002 on a sample of 566 patients had concluded that the Genetic variants of the monoamine oxidase A (MAOA) have been associated with aggression-, anxiety-, and addiction-related behavior (5). Nevertheless, both main and interactive effects of the MAOA VNTR have been associated with ASPD and its symptomatology. It was noted that people with antisocial personality disorder usually manifest themselves as being irritable and aggressive, engaging in behavior that results in a criminal arrest,and they lack remorse for their actions(6).

Recent experimental work and behavior research has linked individual variation in a functional polymorphism of the monoamine oxidase-A gene (MAOA) to the occurrence of anger-driven aggression (7–10).

Furthermore, there is extensive research examining the relation between a polymorphism in the promoter region of the MAOA gene and antisocial phenotypes, as it has been found that 2-repeat allele-may has effects in modulating aggressive behaviors(11, 12). Historically the clinical condition caused by MAOA deficiency, namely Brunner syndrome, was one of the first disorders to be described in relation to MAOA gene and it is characterized by overt antisocial and aggressive conduct (13). studies have found that interaction of low MAOA transcription polymorphisms with environmental factors such as early abuse, deprivation and iron deficiency anemia has been found to increase the occurrence of violent antisocial behavior(14, 15).

The MAO is a mitochondrial enzyme. MAO family composed of two isoenzymes, termed A (Genbank Acc. No: BC008064.1) and B (Genbank Acc. No: M69177), which despite a substantial structural overlap, are remarkably different for substrate preference, inhibitor selectivity, regional distribution, and functional role in behavioral regulation (16). The two monoamine oxidase (MAO) enzymes, A and B, preferentially deaminatesmonoamine neurotransmitters, such as serotonin, norepinephrine, and dopamine. The genes encoding for both types are mapped on chromosome Xp11.23 (17).

It has been found that the termination of serotonin (5-hydroxytryptamine, 5-HT) neurotransmission is regulated by its uptake by the 5-HT transporter (5-HTT), as well as its degradation by monoamine oxidase (MAO)-A. MAO-A deficiency and hypo-methylations lead to a wide set of behavioral alterations, including perseverative behaviors,social deficits and depression(18–21). In addition to that cognitive functions like problem solving were found to be affected by mutations in this gene when it is combined with iron deficiency (ID) in the rhesus monkey(14, 22).

Previous evidence of gene-by-environment interactions associated with emotional and behavioral disorders is contradictory. Differences in findings may be related to variation in the dose of the environmental factor, or failure to take account of gene-by-gene interactions (23).

lastly, the studies of the association between MAOAgenes with childhood maltreatment to the development of antisocial behaviors have been replicated, but not consistently. Several methodological issues can justify these discrepancies, including diverse sample, population genetic heterogeneity and exposure to the various adverse environmental factors (24–26).The aim of this study is to analyze the missense single nucleotides polymorphisms in MAOA gene that could result in the occurrence of antisocial and violent behaviors. Up to our knowledge, this will be the first bioinformatics study to address this topic.

## 2. Material and Methods

### 2.1. Retrieving nsSNPs

SNPs associated with MAOA gene were obtained from the Single Nucleotide Polymorphism database (dbSNP) in the National Center for Biotechnology Information (NCBI). (http://www.ncbi.nlm.nih.gov/snp/).

The sequence and natural variants of MAOA protein were obtained from the UniProt database as it considered as the most reliable database for protein sequences. (27) (https://www.uniprot.org/).

### 2.2. Identifying the most damaging nsSNPs and disease related mutations

A total number of 342 nonsynonymous Single Nucleotide Polymorphisms (ns SNPs) were found from the NCBI database, all of them were subjected to in silico analysis using nine different algorithms and softwares; SIFT, PROVEAN, PolyPhen-2, SNPs&GO, PhD-SNP, PMUT, I-mutant, GeneMANIA and Chimera.

#### 2.2.1. SIFT Server

Phenotypic effects of amino acid substitution on protein function were predicted by using Sorting Intolerant From Tolerant (SIFT) server, which is a powerful tool that uses sequence homology. A list of nsSNPs (rsIDs) from NCBI’s dbSNP database were submitted along with the (original) sequence to SIFT to predict tolerated and deleterious substitutions for every position of a protein sequence. The server divides the results into “Deleterious” and “Tolerated”, nsSNPs with SIFT score ≤ 0.05 were classified as deleterious and are further analyzed to identify the damaging ones, and those > 0.05 were classified as tolerated and are not further analyzed. The server provides fast results but can not assess the result for SNPs that share the same position at the same time. (Available at: http://sift.bii.a-star.edu.sg/)(28–31).

#### 2.2.2. Provean Server

(Protein Variation Effect Analyzer) is the second software tool used. It predicts the effect of an amino acid substitution on the biological function of a protein. It can predict single amino acid substitutions, in-frame insertions, deletions, and multiple amino acid substitutions. The results are obtained as either “Deleterious” if the prediction score was ≤-2.5, while score > −2.5 indicates that the variant is predicted to have a “Neutral” effect. (Available at: http://provean.jcvi.org/index.php)(32, 33).

#### 2.2.3. Polyphen-2 Server

Polymorphism Phenotyping v2.0 (PolyPhen-2) is another online tool that predicts the possible effects of an amino acid substitution on the structure and function of the protein by analysis of multiple sequence alignment. The results are classified into “probably damaging” that is the most disease causing with a score near to 1 (0.7-1), “possibly damaging” with a less disease causing ability with a score of 0.5-0.8 and “benign” which does not alter protein functions with a score closer to zero; (0-0.4). We used a patch query as it provide fast and accurate way to get the result. (Available at: http://genetics.bwh.harvard.edu/pph2/)(27, 34, 35).

#### 2.2.4. SNAP2

It is another tool to predict the Functional effects of mutations. SNAP2 is a trained classifier that is based on a machine learning device called “neural network”. After entering the protein FASTA sequence as an input query, It can distinguish between effect and neutral variants/non-synonymous SNPs by taking a variety of sequences and variant features into account. The most important input signal for the prediction is the evolutionary information taken from an automatically generated multiple sequence alignment. Also structural features such as predicted secondary structure and solvent accessibility are considered. Although it takes a longer time than the previous software to get the result, it has a significant accuracy level (a sustained two-state accuracy (effect/neutral) of 82%)(36). It is available at https://rostlab.org/services/snap2web/.

#### 2.2.4. SNPs & Go server

An online web server that used to ensure the disease relationship with the studied single nucleotide polymorphisms SNPs. It gives three different results based on three different analytical algorithms; Panther result, PHD-SNP result, SNPs&GO result. Each one of these results is composed of three parts, the prediction which decides whether the mutation is neutral or disease related, reliability index (RI), and disease probability (if >0.5 mutations are considered as disease causing nsSNP). (Available at: http://snps-and-go.biocomp.unibo.it/snps-and-go/) (37).

#### 2.2.5. PMUT Server

Is a neural network dependent web-based tool used for the prediction of pathological variants on proteins. The prediction results are classified as “Neutral” or “Disease”. It is very fast and accurate, It is available at (http://mmb.irbbarcelona.org/PMut) (38).

### 2.3. GeneMANIA

A user-friendly web interface tool designed to asses protein function, analyzing submitted gene lists and prioritizing genes for functional assays. It provides lists with functionally similar genes that it identifies and recognizes using available genomics and proteomics data. It is an accurate method to study gene –gene interaction. (Available at: (http://www.genemania.org/)(39).

### 2.4. Modeling nsSNP locations on protein structure

#### 2.4.1 Hope project

It is a webserver that used to analyze the effect of single point mutation on the structural level of protein. it best way to visualize the mutation since it is to create a report consist of figures,animation, 3d structure and mutation report just by submitting the protein sequence and mutation (40). It is available at http://www.cmbi.ru.nl/hope/.

#### 2.4.2 Chimera

Chimera 1.8 software is used for visualization and editing of the three dimensional structure of the protein. It visualizes the original amino acid with the mutated one to see the impact that can be produced. It is a highly extensible program for interactive visualization and analysis of molecular structures and related data, including density maps, supramolecular assemblies,trajectories, and conformational ensembles. High-quality images and animations can be generated. (Availabe at: http://www.cgl.ucsf.edu/chimera)(41).

## 3. ConSurf server

It is a web server that suggests evolutionary conservation reviews for proteins of known structure in the PDB. ConSurf detects similar amino acid sequences and runs multi alignment approaches. The conserved amino acid across species detects its position using specific algorisms (42).

## 4. Variant Effect Predictor (VEP)

The Ensembl Variant Effect Predictor software provides toolsets for an organized approach to annotate and assist for the prioritization of mutations. The input data format was a list of variant identifiers while the output was filtered by choosing 1000 genome combined population to expend the population coverage (43).

## 5. Result

Our results of this study were deduced from analysis of 566 SNPs related to MAOA gene which downloaded from NCBI, 174 SNPs of them were observed as missense mutation. These SNPs were further analyzed by functional and structural softwares to detect the disease associated mutant.

38 SNPs out of 174 SNPs were predicted to be (affect, deleterious, probably damaging, effect); after anlysed by SIFT, PolyPhen-2, PROVEAN, SNAP2 soft wares which used to detect the fuctional and structural change on the protein.(Table 1). In addition 6 SNPs out of the 38 SNPs prognosticated to be (disease)when analysed by SNP&GO, PHD and PMut softwares; it function to ensure disease relationship(Table 2). Eventually, 18 variants were branched from the 6 disease related SNPs by VEP tool. (Table 3).

**Table.1.**
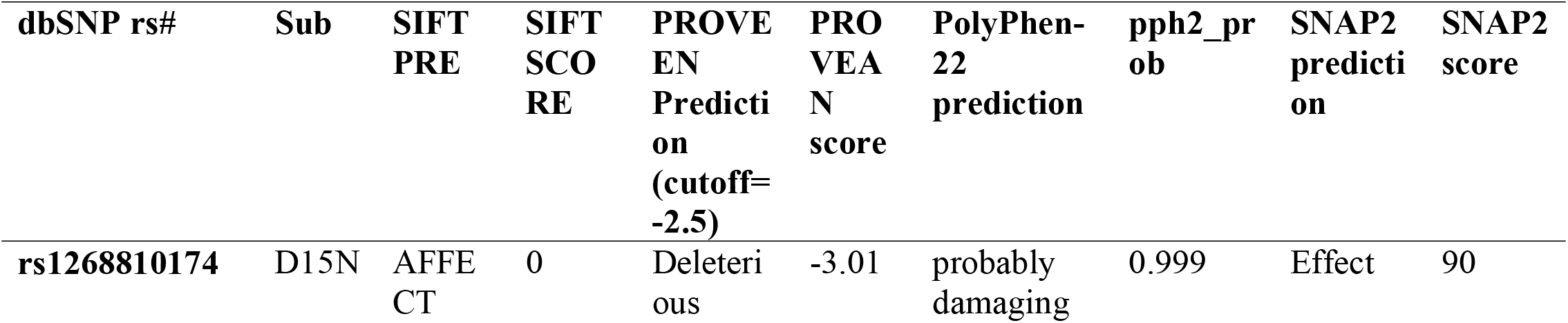
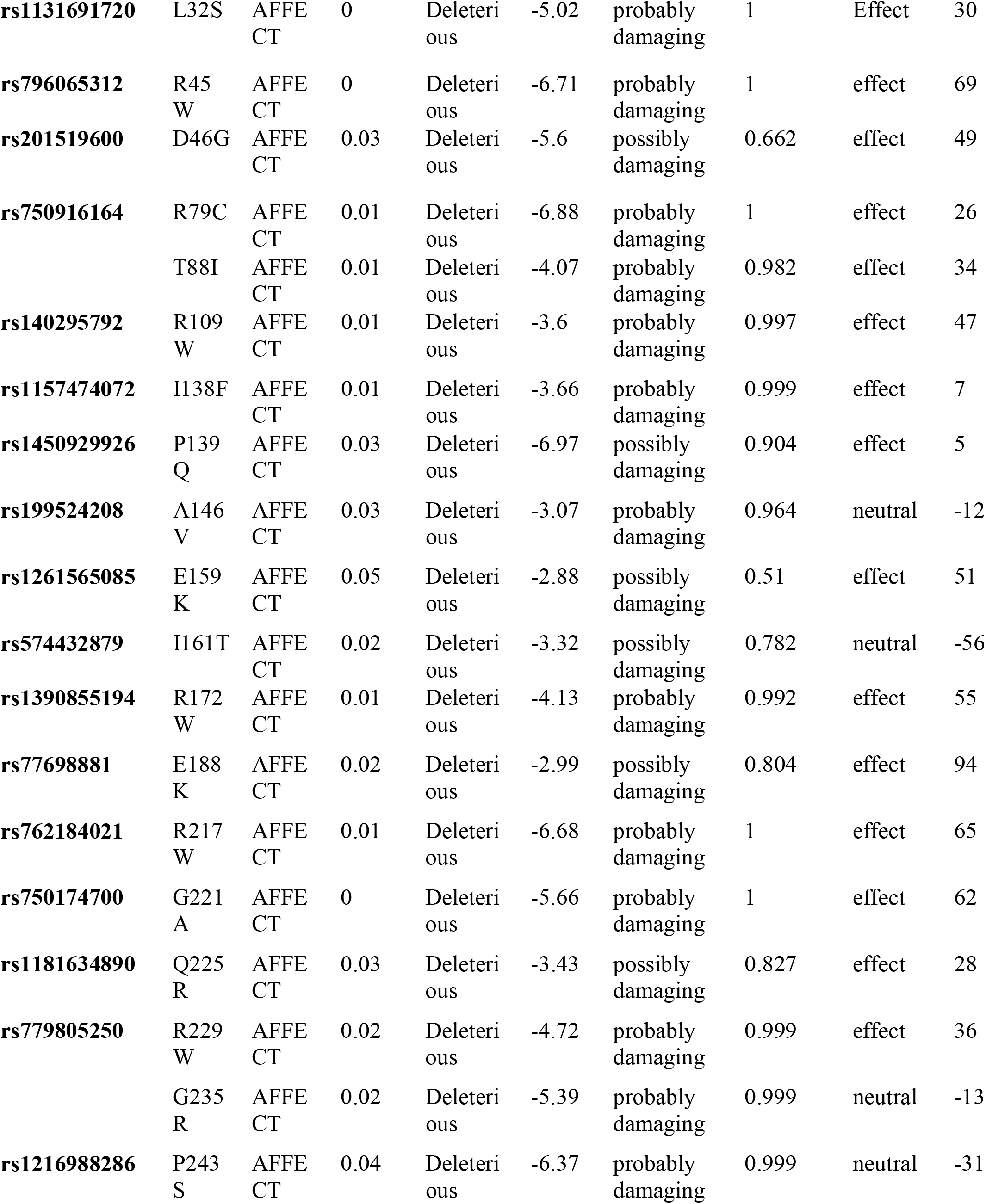
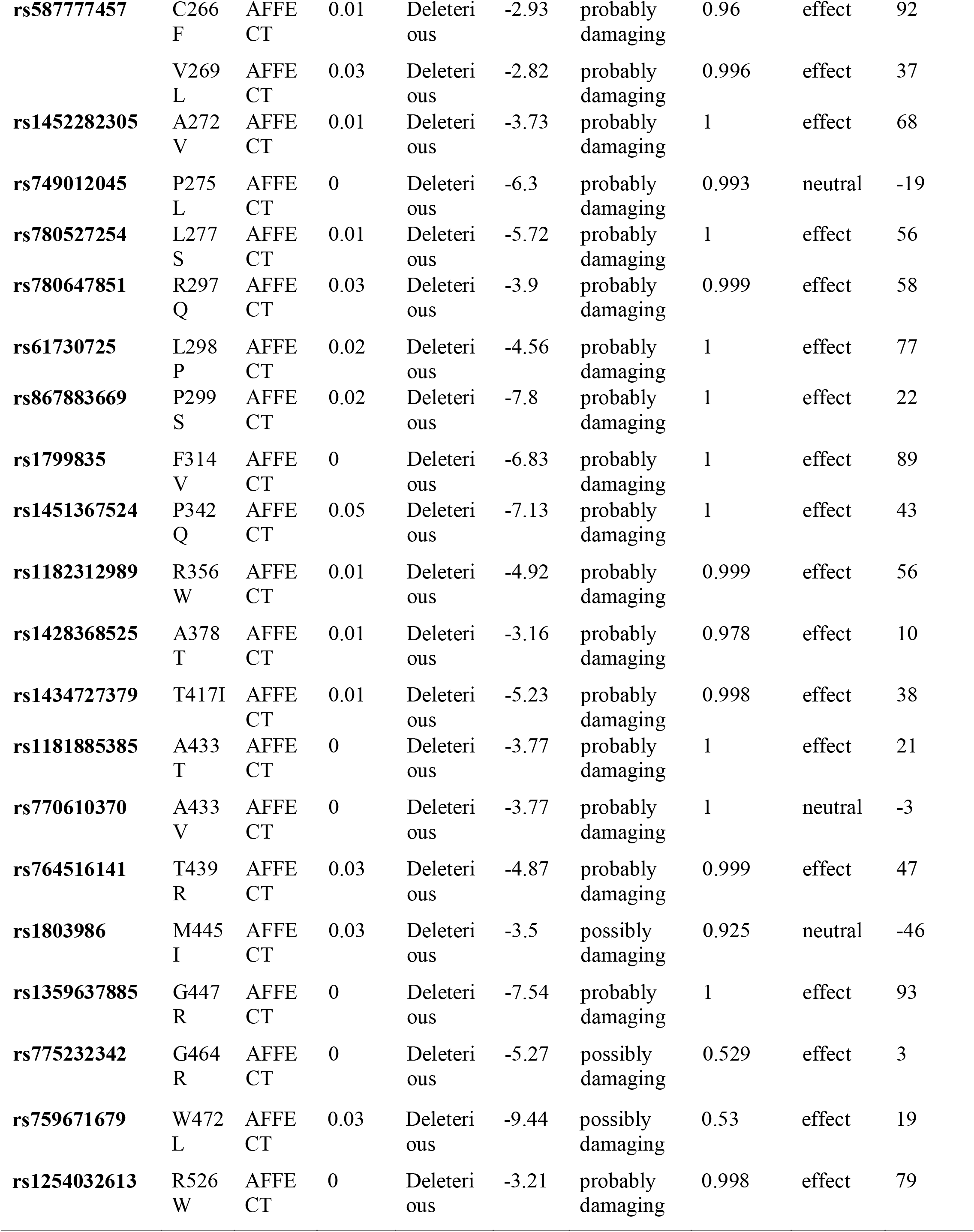
Deleterious nsSNPs associated variations predicted by SIFT, PolyPhen-2, PROVEAN, SNAP2 soft wares.

**Table.2.** Disease effect nsSNPs associated variations predicted by Pmut, SNP&GO, PHD and PMut softwares.

| dbSNP rs# | sub | Prediction phd | R I | Probability | Prediction snp and go | R I | Probability | Pmut prediction | Pmut score |
| --- | --- | --- | --- | --- | --- | --- | --- | --- | --- |
| **rs1131691720** | L32S | Disease | 7 | 0.87 | Disease | 4 | 0.725 | Disease | 0.90<br>(93%) |
| **rs796065312** | R45W | Disease | 8 | 0.921 | Disease | 6 | 0.816 | Disease | 0.90<br>(93%) |
| **rs750174700** | G221<br>A | Disease | 4 | 0.679 | Disease | 2 | 0.581 | Disease | 0.72<br>(86%) |
| **rs1799835** | F314<br>V | Disease | 8 | 0.886 | Disease | 6 | 0.781 | Disease | 0.75<br>(88%) |
| **rs1181885385** | A433<br>T | Disease | 4 | 0.717 | Disease | 1 | 0.527 | Disease | 0.83<br>(90%) |
| **rs1359637885** | G447<br>R | Disease | 5 | 0.756 | Disease | 4 | 0.717 | Disease | 0.92<br>(94%) |

**Table (3).**
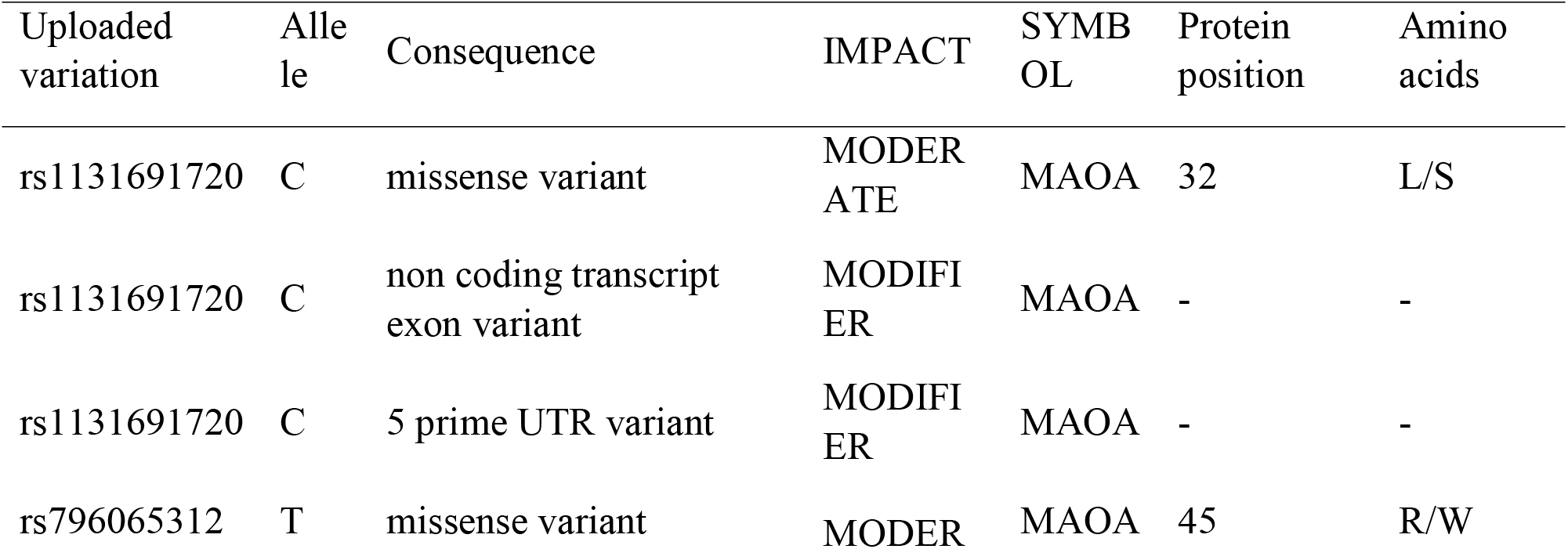
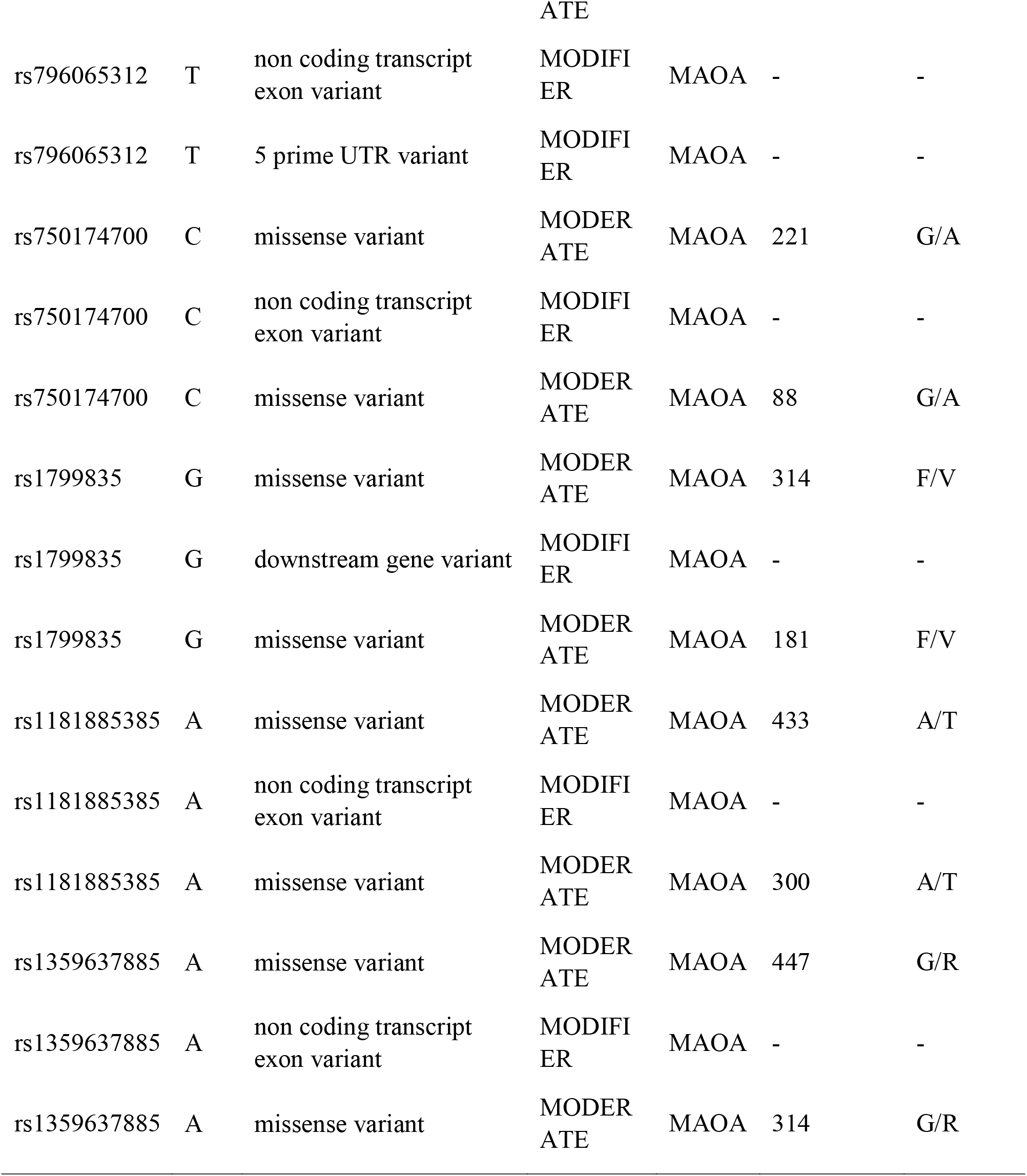
Shows variant consequences, transcripts, and regulatory features by VEP tool:

The reference sequence was submitted to RaptorX, to get the structural prediction of the protein, which was further displayed by Chimera software; to visualize the changes in the wild and mutant amino acids.(Figure 1–6). Then gene to gene relations was identified using geneMAINA software (Figure7).moreover, the conservation of the SNPs among different species was detected using ConSurf software (Figure8).

**Figure1:**
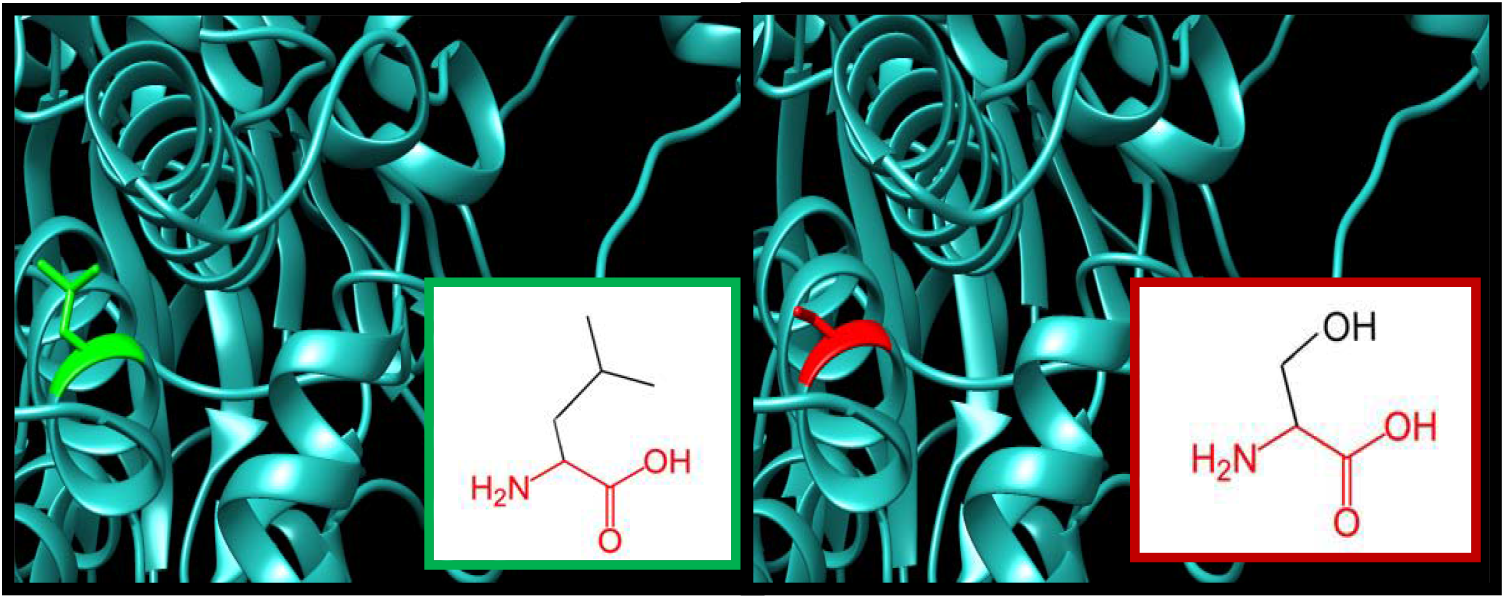
L32S: Leucine (green color) changed to Serine (red color) at position 32. The green color represents wild amino acid while the red color represents mutant amino acid and the cyan color represents the background structure of the MAOA protein.

**Figure 2:**
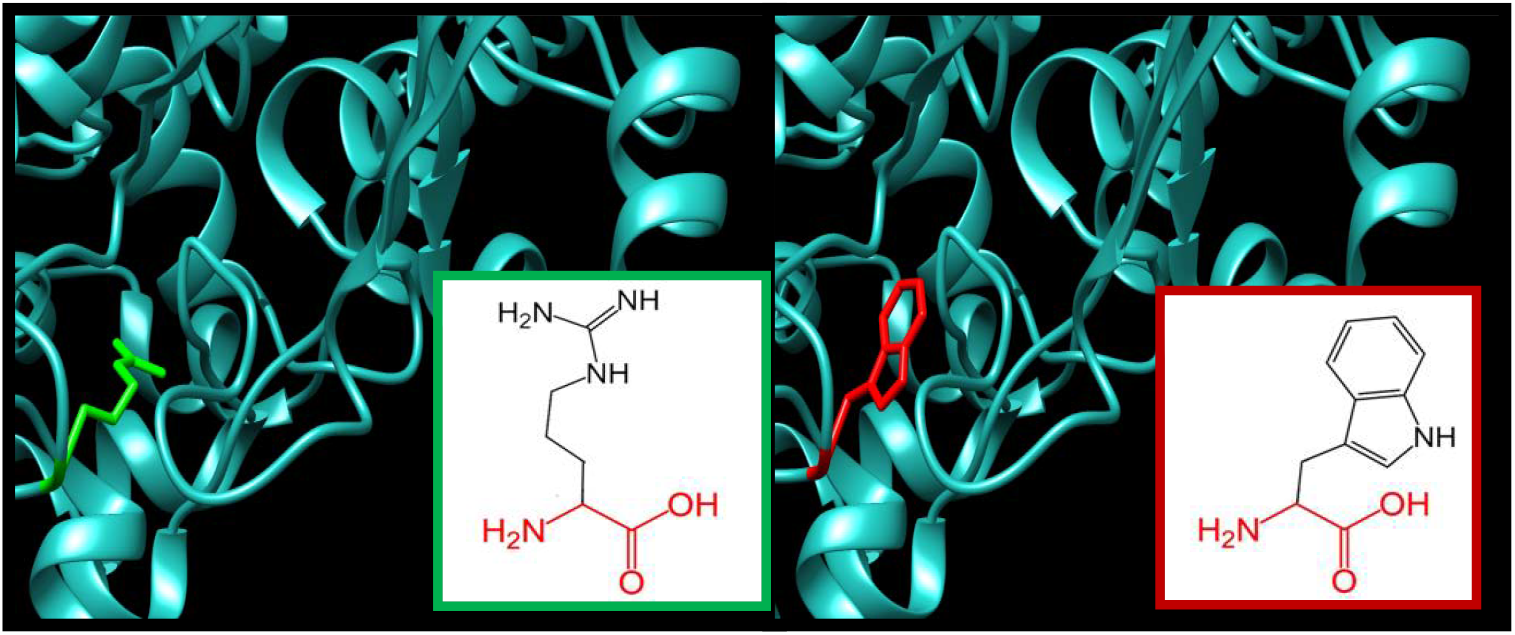
A45W: Arginine(green color) changed to Tryptophan (red color) at position 45. The green color represents wild amino acid while the red color represents mutant amino acid and the cyan color represents the background structure of the MAOA protein.

**Figure3:**
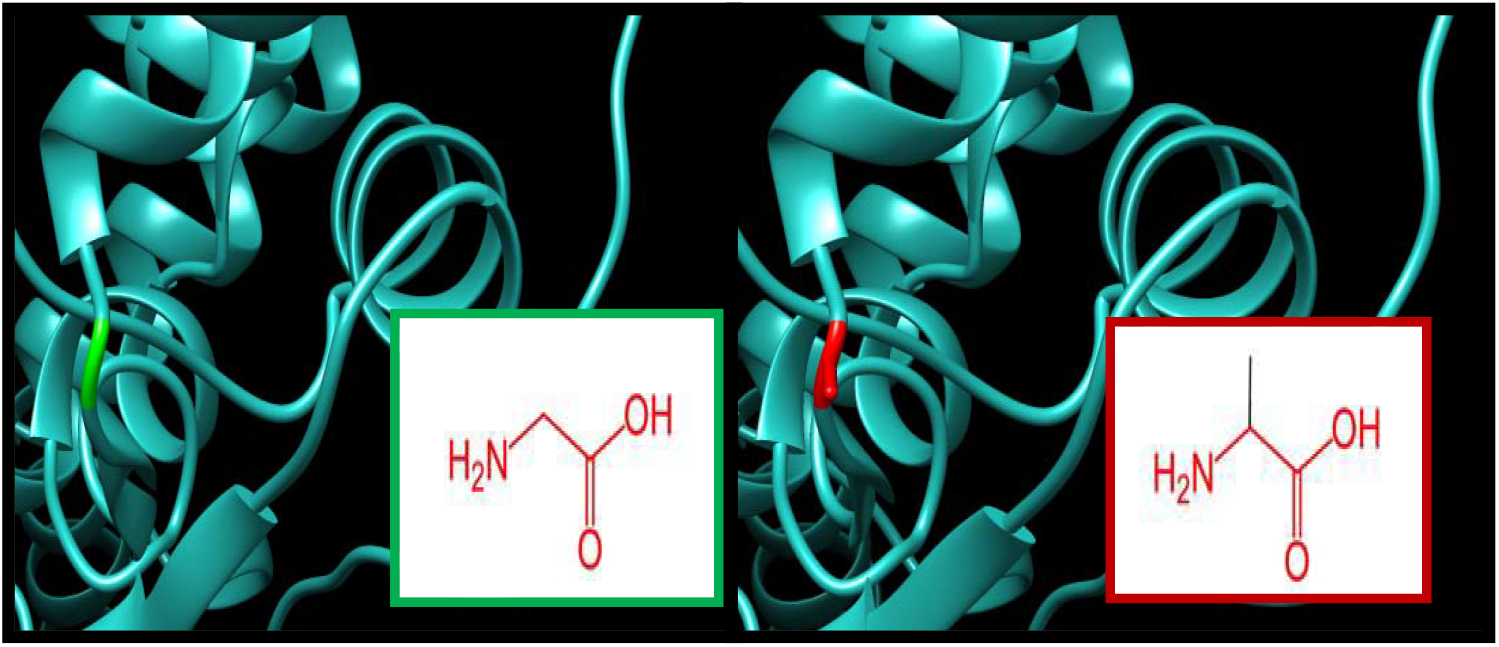
G221A: Glycine (green color) changed to Alanine(red color) at position 221.The green color represents wild amino acid while The red color represents mutant amino acid and the cyan color represents the background structure of the MAOA protein.

**Figure4:**
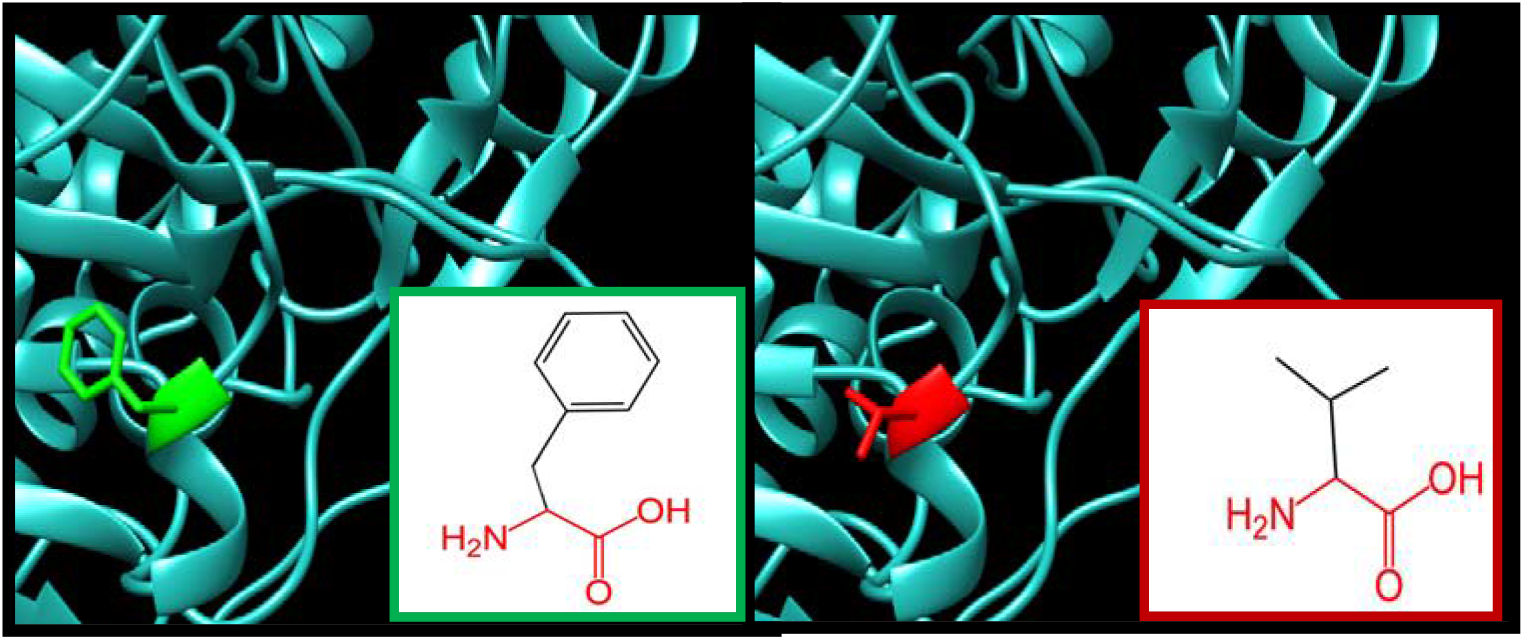
F314V: Phenylalanine (green color) changed to Valine(red color) at position 314. The green color represents wild amino acid while The red color represents mutant amino acid and the cyan color represents the background structure of the MAOA protein.

**Figure 5:**
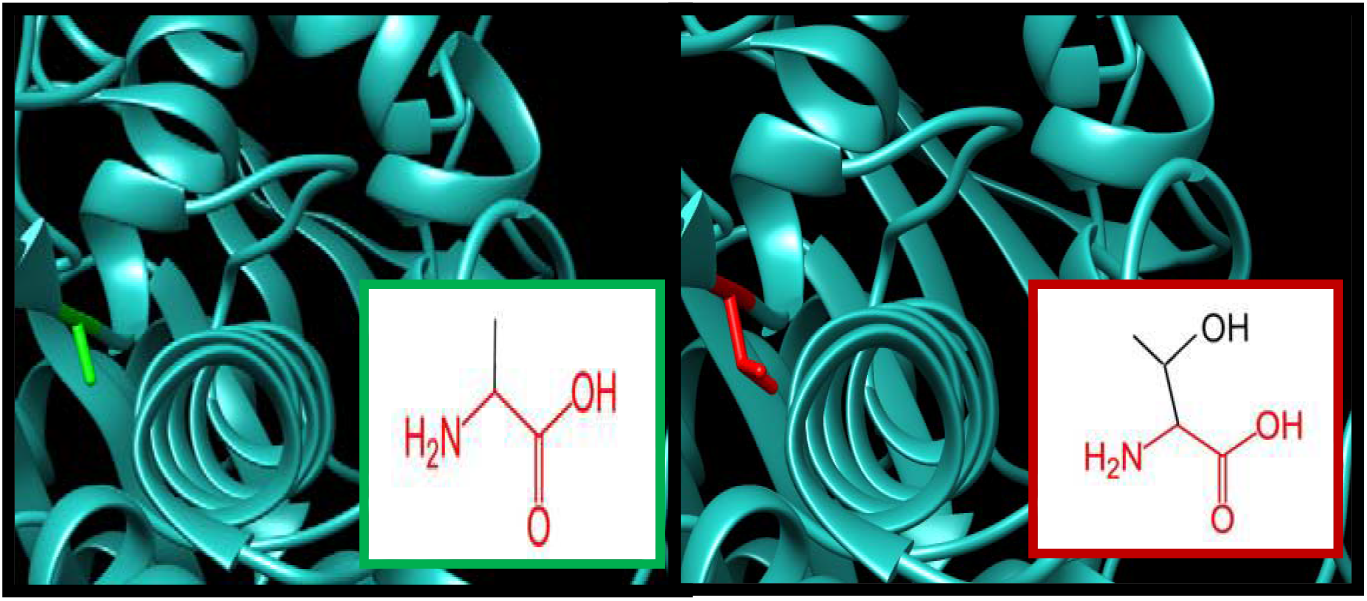
A422T: Alanine(green color) changed to Threonine(red color) at position 433. The green color represents wild amino acid while The red color represents mutant amino acid and the cyan color represents the background structure of the MAOA protein.

**Figure6:**
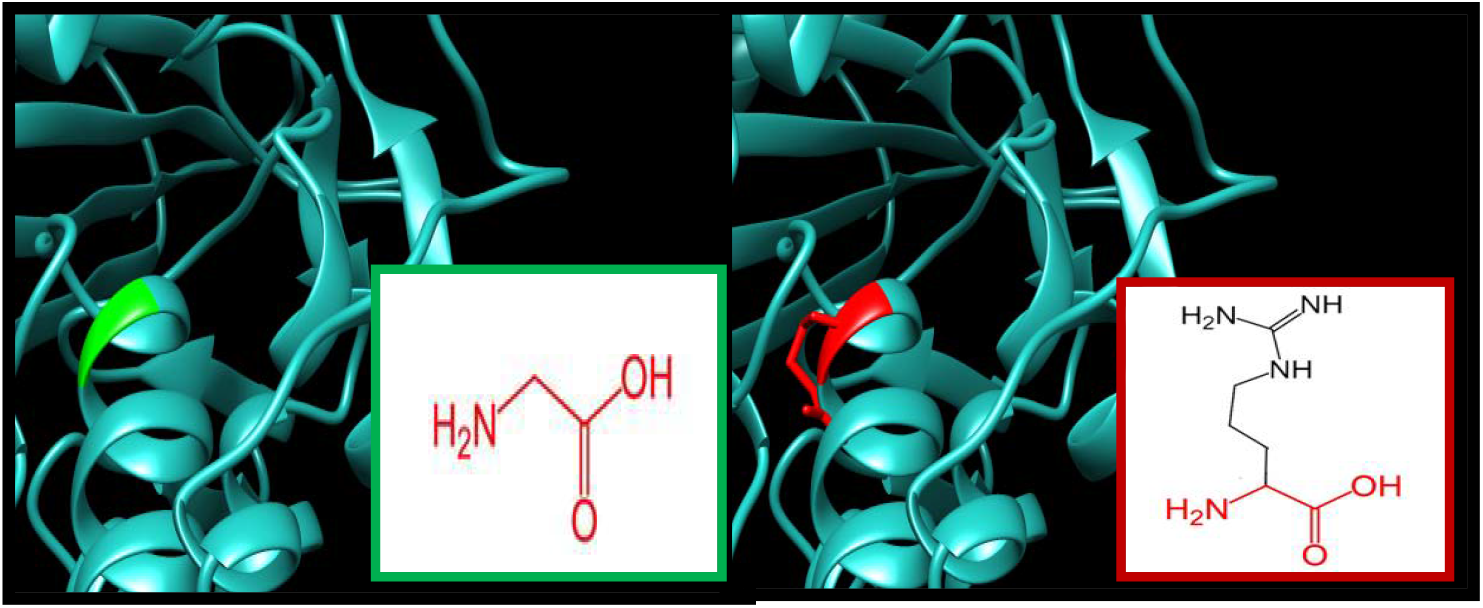
G447A: Glycine (green color) changed to Arginine (red color) at position 447. The green color represents wild amino acid while The red color represents mutant amino acid and the cyan color represents the background structure of the MAOA protein.

**Figure(7):**
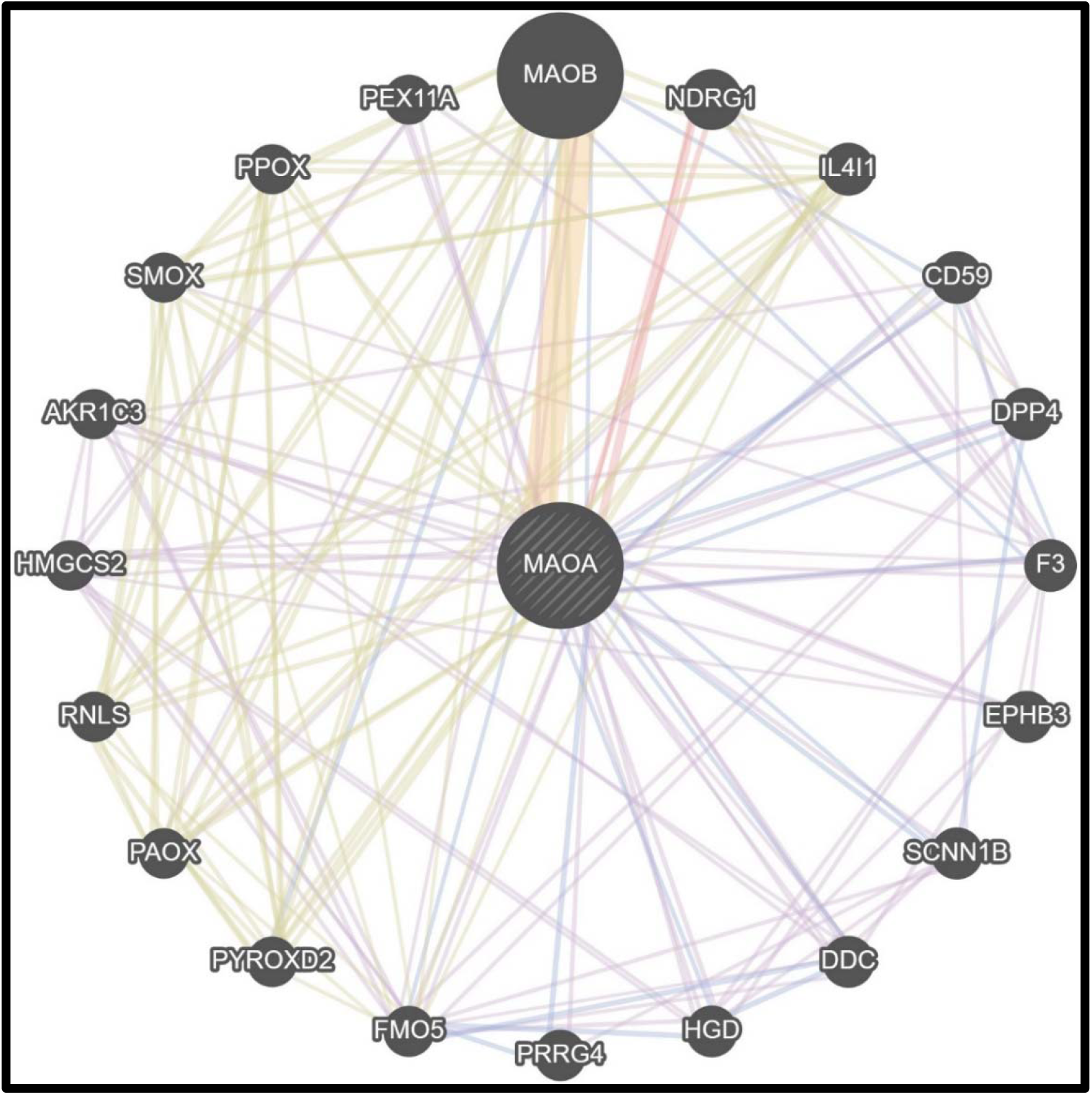
shows gene-gene interaction by geneMANIA. The lines represent the complicated interactions between MAOA gene and other genes.

**Figure(8).**
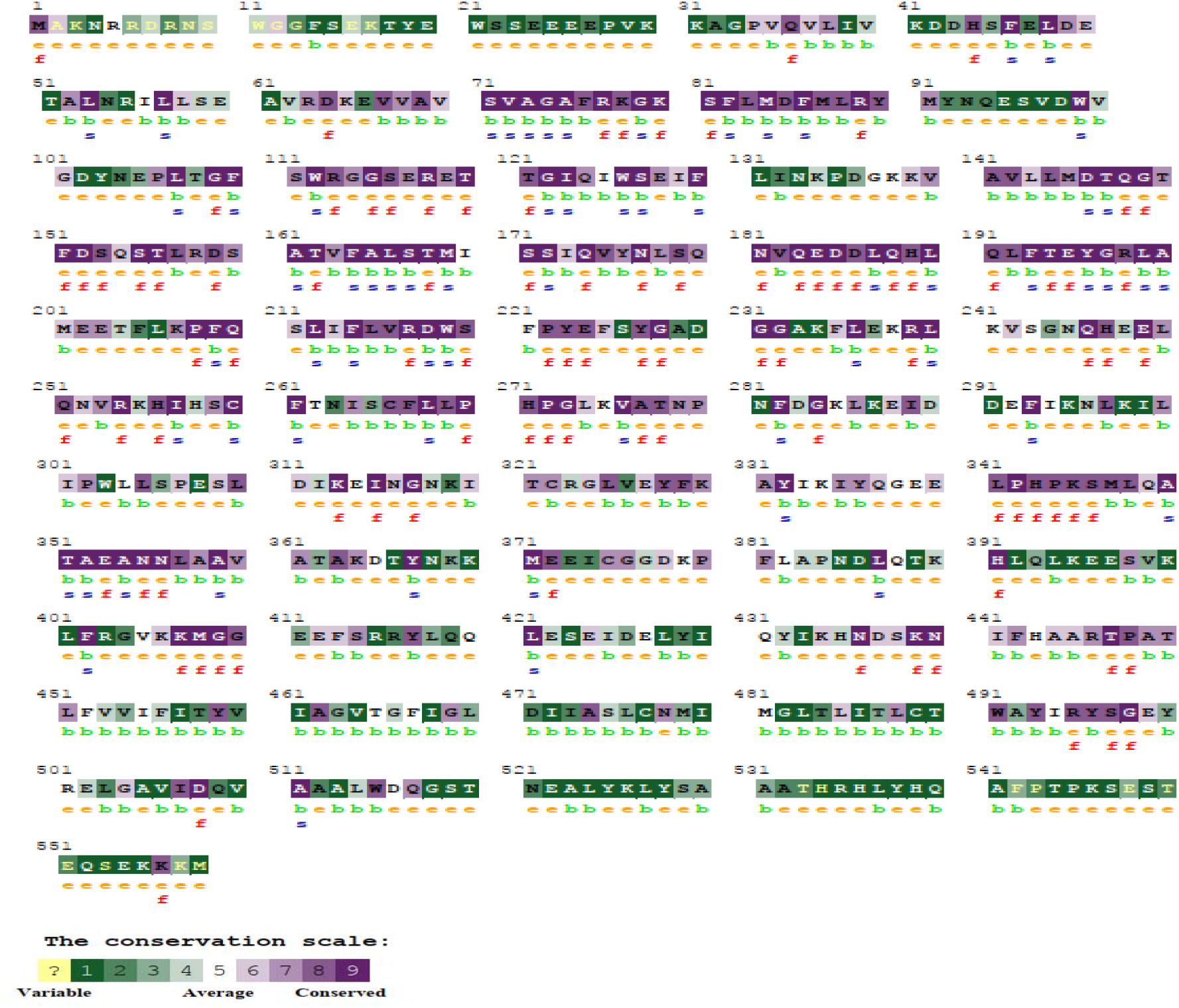
The conserved amino acids across species in MAOA protein were determined using ConSurf. e: exposed residues according to the neural-network algorithm are indicated in orange letters. b: residues predicted to be buried are demonstrated via green letters. f: predicted functional residues (highly conserved and exposed) are indicated with red letters. s: predicted structural residues (highly conserved and buried) are demonstrated in blue letters. I: insufficient data (the calculation for this site performed in less than 10% of the sequences) is demonstrated in yellow letters.

## 6- Discussion

Six novel missense SNPs were found in MAOA gene to effect protein function and structure. Out of 566 SNPS downloaded from NCBI, 34 SNPs have been found to be damaging by four functional software. Which were further analyzed by SNP&GO and PMut, 6 SNPs were found to be damaging, as illustrated in table (1 and 2).

Project hope was used to study the physiochemical changes inflicted by the missense mutations. All the SNPs were found to be located at Fad/Nad(P)-Binding Domain Superfamily IPR036188 and Flavin Amine Oxidase IPR001613 domain which indicates this SNPs is likely to be conserved. In all of the SNPs the mutant amino acid was bigger in size than the wild type except in F314V and L32S SNPs. In addition all the mutant amino acids were found to be different in hydrophobicity when compared with the wild type amino acid. All these changes will affect protein function.

Structural changes were further studied using chimera software, where changes in the amino acids Cleary change the structural of the MAOA protein as evident in figure(1–6).

The interaction and function of MAOA gene were studied using gene mania. where the predicted functions were: amine metabolic process, cellular amine metabolic process, cellular biogenic amine metabolic process, cellular response to xenobiotic stimulus, oxidoreductase activity, acting on the CH-NH2 group of donors, oxidoreductase activity, acting on the CH-NH2 group of donors, oxygen as acceptor, response to xenobiotic stimulus and xenobiotic metabolic process. The MAOA gene was found to have interactions with many other genes like CD50,F3,HGD and others as shown in Figure (7).

The Variant Effect Predictor annotates mutations using an extensive array of reference data from previously detected mutations, evidence based results, and estimation of biophysical consequences of mutations; and that is what makes VEP an accurate web based tool..(43) VEP described regulatory consequences for several mutations, including 10 mutations within the coding region, 5 mutations within a non-coding region. briefly, mutations within coding region affect the protein function, while mutations within non-coding regions can significantly affect disease and could contribute in the phenotypic feature and RNA-binding proteins (RBPs) (44, 45), while mutations in the upstream, downstream, 5’-, and 3’-UTRs might affect transcription or translation process (46). The consequences are shown in Table(3).

We also used ConSurf web server; the nsSNPs that are located in highly conserved regions and predicted to cause structural and functional impacts on MAOA protein Figure (8)

Previous researches show that regulatory variations in MAOA to moderate effects of childhood maltreatment on male antisocial behaviors (26). while another research showed that certain SNPs polymorphism (1460 C>T) is associated with other conditions leading to the development of depressive symptoms in menopausal women (47). On the other hand another research underwent in Chinese population, the interaction between catechol-O-methyltransferase (COMT), Ala22/72Ser and MAOA T941G polymorphisms, and stressful life events (SLE) in the academic pressure and the relation to aggressive behavior development were also identified (48). This SNP was not available in the data we acquired from NCBI.Furthermore, these studies do indicate the significant role of MAOA gene polymorphisms in aggrieve behavior development.

There are other studies that assess the contribution of gene mutations in the mediation of directed aggression in humans, as in the study done on a population of 328 German subjects which reported that, the functional polymorphism in Chatechol-O-methyltranferese (COMT) gene can alter the phenotype of suicidal and anger behaviors(49).

From the previous evidence we believe that a screening for MAOA, COMT and other similar genes mutation in suspected individuals may help in detecting earlier violent behavior in popullations which will provide the chance for earlier intervention, leading to decrease in the incidence of aggressive behavior and criminal act.

As we report in this paper further illustrate the significant role that genetics plays in the development of violent behavior, we still believe that every one has a choice at some point in his life whether to succump to his inner devil or not.

This study is limited, as it does not cover the environmental factor important for the development of antisocial behavior, nevertheless this study will lay the foundation for further clinical research in this phenomenon.

## 7- Conclusion

Six novel missense SNPs were found in MAOA gene that affect MAOA protein function and structure, which could predispose to violent and aggressive behavior. Therefore this gene along with other similar genes colud be used as a surveillance tool for early detection of aggressive behavior in the population.

## 9- Acknowledgment

The authors acknowledge the Deanship of Scientific Research at the University of Bahri for supportive cooperation.

## Data Availability

All data are available within this manuscript.

